# On the validity and errors of the pseudo-first-order kinetics in ligand–receptor binding

**DOI:** 10.1101/051136

**Authors:** Wylie Stroberg, Santiago Schnell

**Author notes:** Corresponding author. Email addresses (Wylie Stroberg), (Santiago Schnell).

## Abstract

The simple bimolecular ligand–receptor binding interaction is often linearized by assuming pseudo-first-order kinetics when one species is present in excess. Here, a phase-plane analysis allows the derivation of a new condition for the validity of pseudo-first-order kinetics that is independent of the initial receptor concentration. The validity of the derived condition is analyzed from two viewpoints. In the first, time courses of the exact and approximate solutions to the ligand–receptor rate equations are compared when all rate constants are known. The second viewpoint assess the validity through the error induced when the approximate equation is used to estimate kinetic constants from data. Although these two interpretations of validity are often assumed to coincide, we show that they are distinct, and that large errors are possible in estimated kinetic constants, even when the linearized and exact rate equations provide nearly identical solutions.

## 1. Introduction

In biochemical kinetics, simplifying assumptions that decouple or reduce the order of rate equations for complex reaction mechanisms are ubiquitous. Aside from making theoretical analysis of complex reactions more tractable, order-reducing approximations can greatly simplify the interpretation of experimental data [1, 2]. Experiments performed under conditions that allow for linearization have historically been the preferred method for estimating equilibrium and rate constants because they allow for the isolation of a subset of the interactions [3, 4, 5]. For this reason, when designing an experiment, it is essential to know the necessary conditions for the simplifying assumptions to be valid. Significant theoretical work has been directed at deriving rigorous bounds for the validity of simplifying assumptions [6, 7, 8, 9, 10, 11], but this work often overlooks the manner in which the reduced models are used to interpret experimental results. In many cases, the simplified models are used to estimate equilibrium and rate constants from experimental data [12, 13, 14, for example]. Rarely is the validity of a simplifying assumption analyzed with this utility in mind. To examine how this viewpoint can affect the conditions for validity, we consider the simplest model for ligand–receptor binding with 1:1 stoichiometry [15].

In the simplest case, the binding of a ligand *L* to a receptor *R* is a bi-molecular reversible association reaction with 1:1 stroichiometry yielding a ligand–receptor intermediate complex *C:*

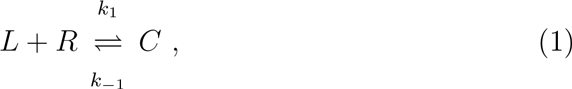

where *k*_1_ and *k*_−1_ are, respectively, the association and dissociation rate constants of the ligand–receptor complex. This reaction scheme is mathematically described by a system of coupled nonlinear second-order differential equations. By applying the law of mass action to reaction (1), we obtain

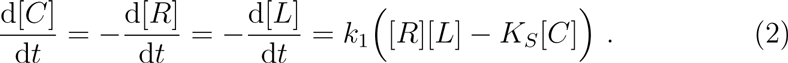

In this system the parameter *K*_*S*_ = *k*_−1_/*k*_1_ is the equilibrium constant [15, 4] and the square brackets denote concentration. Since no catalytic processes are involved, the reaction is subject to the following conservation laws:

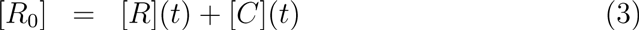

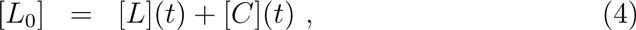

where [*R*_0_] and [*L*_0_] are the initial receptor and initial ligand concentrations. If the bimolecular reaction (1) is initiated far from the equilibrium and in the absence of ligand–receptor complex, the system (2) has the initial conditions at *t* = 0:

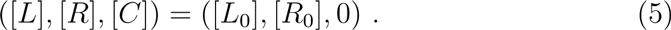

We have expressed quantities in terms of concentration of species. These equations are frequently given in terms of binding site number, using the identity [15]

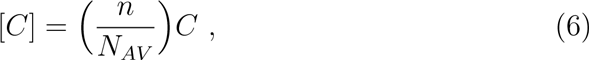

where *n* is the cell density, *N*_*AV*_ is Avogadro’s number, and *C* denotes the number of ligand–bound receptors per cell. We use the concentration formulation here for clarity and without loss of generality.

The system (2) can be solved, subject to the conservation laws [16]. Substituting (3) and (4) into (2) we obtain:

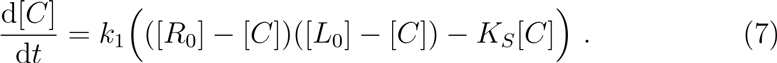

We can rewrite this expression by factoring as follows:

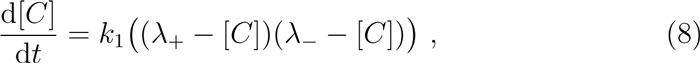

where

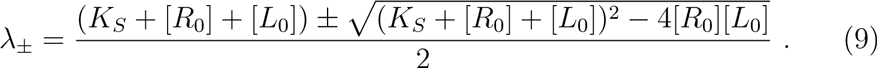

This ordinary differential equation is readily solved subject to the initial conditions (5) as

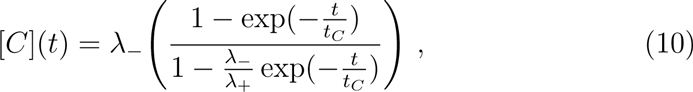

with

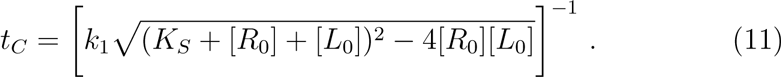

The quantity *t*_*C*_ is the timescale for significant change in [*C*]. In this particular case, *t*_*C*_ can be considered as the time required for the reaction to reach steady-state. Solutions for [*R*](*t*) and [*L*](*t*) can now be constructed by substituting (10) into conservation laws (3) and (4).

Although there is a closed form solution for the reacting species of the simple bimolecular ligand–receptor interaction, experimental biochemists prefer to determine the kinetic parameters of the ligand–receptor binding using graphical methods [15]. One of the graphical methods commonly used consists of plotting the solution of the ligand association assuming no ligand depletion on a logarithmic scale with respect to time. Both the association and dissociation rate constants can be determined using this linear graphical method [17]. Similarly, if one seeks to avoid inaccuracies due to logarithmic fitting, nonlinear regression can be used to fit the kinetic data to a single exponential. However, the use of both of these methods has the disadvantage of making an assumption with respect to the relative concentrations of ligand and binding sites [16].

In the ligand–receptor interaction with 1:1 stochiometry and no ligand depletion it is generally thought that, if the initial ligand concentration is much higher than the initial receptor concentration, i.e.

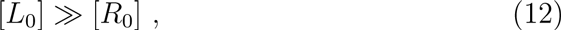

the ligand concentration [*L*] remains effectively constant during the course of the reaction, and only the receptor concentration [*R*] changes appreciably with time [18, 19, 3, 4]. Since kinetic order with respect to time is the same as with respect to [*R*], reaction (1) is said to follow *pseudo-first-order* (PFO) kinetics if the [*R*] dependence is of first order. The rates of second-order reactions in chemistry are frequently studied within PFO kinetics [20, 21]. In the present case, the second-order reaction (1) becomes mathematically equivalent to a first-order reaction, reducing to

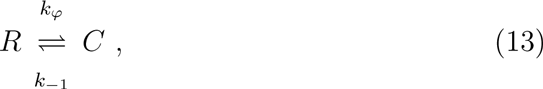

where *k*_*φ*_ ≡ *k*_*1*_[*L*_0_] is the pseudo rate constant. This procedure is also known as the method of flooding [5]. The solution of the governing equations for a reaction linearized by PFO kinetics (or flooding) is straightforward, and is widely employed to characterize kinetics and fit parameters with the aid of progress curves. An error is however present due to the fact that, in actuality, the concentration of the excess reactant does not remain constant [20].

In 1961, Silicio and Peterson [20] made numerical estimates for the fractional error in the observed PFO constant for second-order reactions. They found that the the fractional error is less than 10% if the reactant concentration ratio, [*R*_0_]: [*L*_0_] say, is tenfold. On the other hand, Corbett [22] found that simplified expressions with the PFO kinetics can yield more accurate data than is generally realized, even if only a twofold excess of one the reactants is employed. For ligand–receptor dynamics, Weiland and Molinoff [16] clai that the PFO simplification is acceptable if experimental conditions are such that less than 10% of the ligand is bound. These results indicate that the conditions whereby a second-order ligand–receptor reaction is reduced to first order remain to be well-established.

It is widely believed that second-order reactions can be studied by PFO kinetics using progress curves only when the excess concentration of one of the reactants is large [21, 5, for example]. However, contrary to the widely established knowledge, Schnell and Mendoza [10] have found that the condition for the validity of the PFO in the single substrate, single enzyme reaction does not require an excess concentration of one of the reactant with respect to the other. In the present work, we derive the conditions for the validity of the PFO approximation in the simple ligand–receptor interaction. Additionally, we show two fundamentally different methods of assessing the validity of the approximation. The first compares the exact and approximate solutions to the rate equations under identical conditions. The second measures the veracity of parameters estimated by fitting the approximate model to data. Although these two measures of validity are generally assumed to coincide, we show that they are quantitatively and qualitatively distinct. In Section 2 the reduction of the ligand–receptor association by PFO kinetics is summarized followed by its dynamical analysis in Section 3. The new validity condition is derived in Section 4, and an analysis of the errors observed with the PFO kinetics is presented in Section 5. This is followed by a brief discussion (Section 6).

## 2. The governing equations of the ligand–receptor dynamics with no ligand depletion

In ligand–receptor dynamics with 1:1 stochiometry and no ligand depletion, the second-order ligand–receptor interaction in reaction (1) is neglected when condition (12) holds; the reaction effectively becomes first order since the concentration of the reactant in excess is negligibly affected. This is equivalent to assuming that

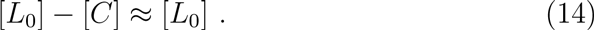

The alternative case, in which the depletion of the receptor is assumed to be negligible, is shown to be symmetric in Appendix A. By substituting (14) into (7), the equation can be simplified as follows:

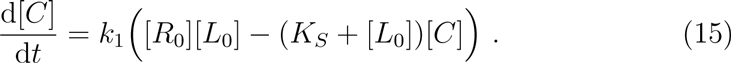

Note that this equation can also be obtained by applying the law of mass action to reaction scheme (13). The solution for (15) with the conservation laws (4) and (5) can be obtained by direct integration [23]:

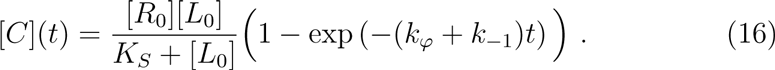

Note that expressions for [*R*](*t*) and [*L*](*t*) can again be obtained by substituting (16) into (3) and (4), respectively.

Experimentally, the clear advantage of applying the pseudo-first-order kinetics to the ligand–receptor reaction is that, as shown by equation (16), it provides solutions that can be linearized by using a logarithmic scale to fit progress curves of the interacting species and thus it could lead to complete reaction characterization, namely the rate constants *k*_1_ and *k*_−1_

As we have previously pointed out, it has been assumed that the condition [*L*_0_] ≫ [*R*_0_] implies

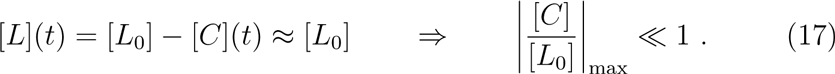

Up to this point, most of the scientists using the PFO kinetics assume that it is reasonable to overestimate the maximum complex concentration when there is a ligand excess, because all receptor molecules could instantaneously combine with ligand molecules, i.e. [*C*]_max_ = [*R*_0_ However, this simplification is unrealistic from the biophysical chemistry point of view as it will assume that in the conservation law (4) all ligand molecules are only in one form: the free ligand [see, equation (17)]. In the next section, we obtain a more a reliable estimate of [*C*]_max_ by studying the geometry of the phase plane of system (2). This will permit us to make a better estimate of the conditions for the validity of the PFO ligand–receptor dynamics (1).

## 3. Phase-plane analysis leads to conditions for the validity of the pseudo-first-order kinetics

The phase-plane trajectories of system (2) are determined by the ratio of d[*C*]/d*t* to d[*L*]/d*t:*

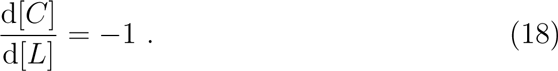

This expression is integrated to obtain the family of solution curves:

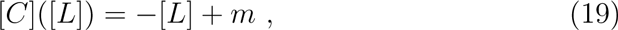

where

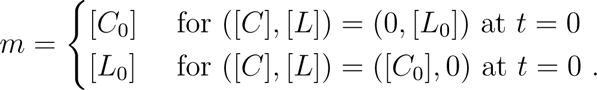

The use of [*L*_0_] as the constant in (19) for initial conditions on the horizontal axis follows from the relation d[*C*]/d*t* =—d[*L*]/d*t*.

The phase plane is divided into two regions by the nullcline

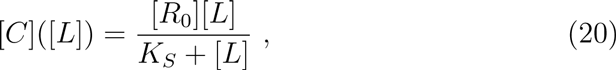

obtained by setting d[*C*]/d*t* = 0 or d[*L*]/d*t* = 0 in (2). Note that for [*L*] = *K*_*S*_ (equilibrium constant), [*C*] = [*R*_0_]/2. The nullcline converges at large ligand concentrations because

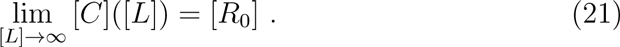

The phase plane trajectories (19) and its nullcline (20) are show in Fig. 1. The trajectory flow is attracted by a unique curve, which is a stable manifold and is equivalent to the nullcline for this case. All trajectories tend to this manifold as they approach the steady state as *t* → ∞ [24].

**Figure 1:**
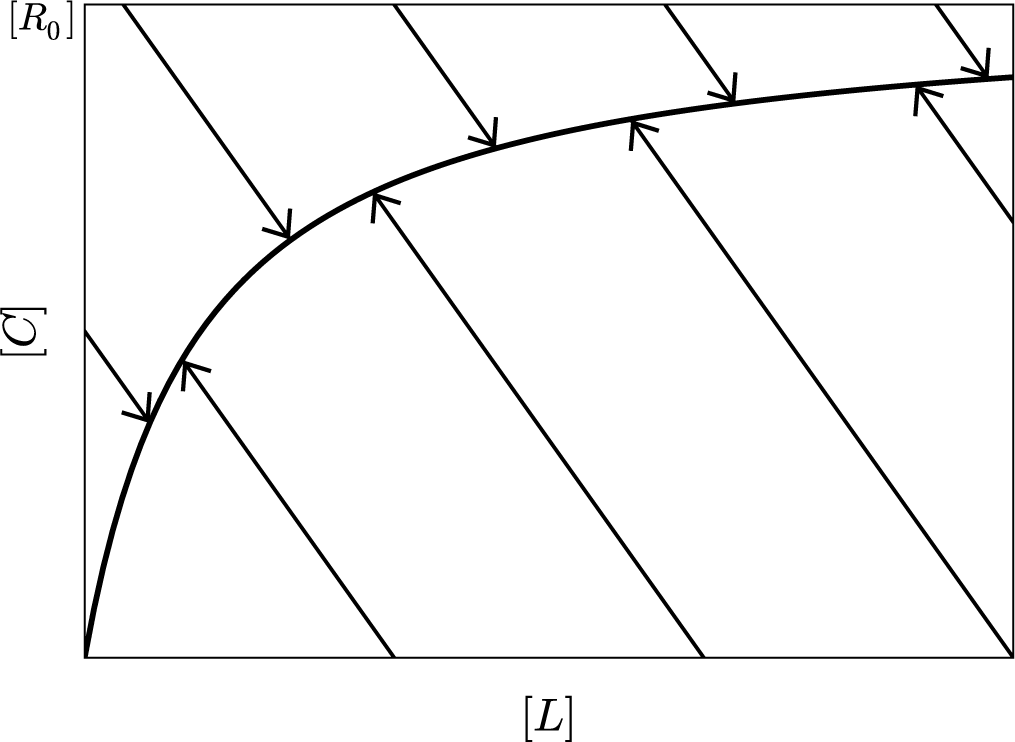
Phase-plane behavior of the ligand receptor reaction (1). The solid curves with arrows are the trajectories in the phase planes, which are described by (19). The trajectories tend to a stable manifold as they approach to the steady-state. In this case, the manifold is the nullcline (20) of the system, which converges to [*R*_0_] for large ligand concentrations.

Binding of ligand to cell surface receptors has been amenable to *in vitro* experimental investigation for the past four decades [25]. In the typical experimental approach, isolated membranes possessing free receptors are studied using ligands as pharmaceutical agents [26]. The reaction mixture is free of ligand–receptor complex at the beginning of the experiment, that is the initial conditions are like those stated in (5). It is important to note that for a ligand–receptor interaction with trajectories departing from the positive horizontal axis, i.e. with initial conditions (5), the trajectories are boundedby

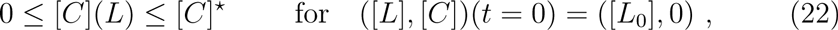

where [*C*]* is the ligand–receptor complex concentration at the steady-state, and is equivalent to the maximum ligand–receptor complex concentration ([*C*]_max_) that the trajectories can reach if they depart from the positive horizontal axis. [*C*]* can be estimated from the intersection of (19) and (20) or by estimating the steady-state value of the ligand–receptor complex concentration in (10), that is

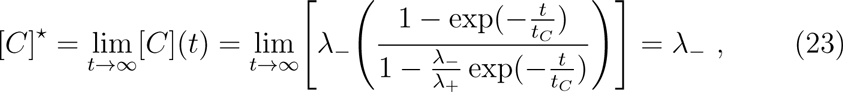

where λ_ is given by (9).

It suffices therefore to investigate the behavior of the ratio of the solution (10) at steady-state to [*L*_0_], which we do in the next section.

## 4. Derivation of a new sufficient condition for the validity of the pseudo-first-order kinetics

To derive a mathematical expression in terms of the kinetic parameters for condition (17), we will use the fact that, for initial conditions given by (22), [*C*]_*max*_ is the concentration given by allowing the reaction described by (10) to go to steady-state. We can now formulate (17) as follows:

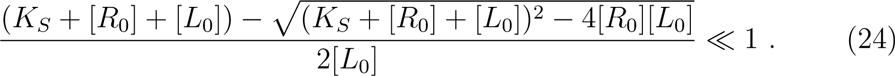

This can be rewritten as

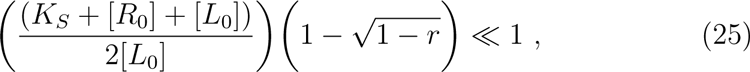

with

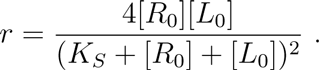

At this point it is convenient to nondimensionalize the above expression by using reduced concentrations. Scaling with respect to *K*_*S*_, equation (25) becomes

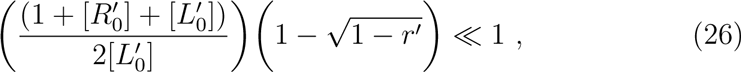

where

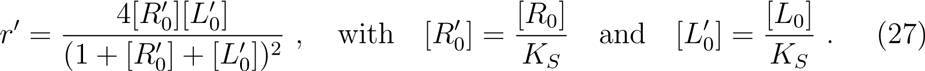

Quadratic expressions similar to (26) are common in chemical kinetics. For practical use in the analysis of chemical kinetics experiments, quadratic expressions are generally replaced with simpler expressions. Noting that

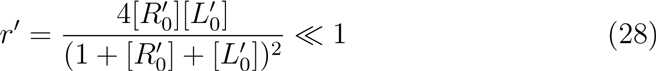

for any value of 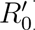 and 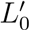 (for more details, see Appendix B), we can then calculate a Taylor series expansion of (26) to obtain right-hand factor of

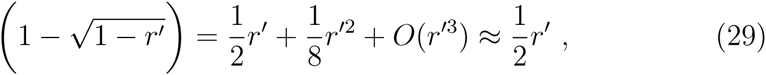

which simplifies (26) to

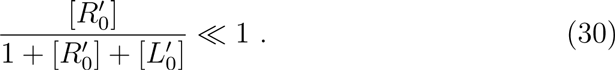

This is a simple analytical expression for the condition for the validity of PFO kinetics. Note that condition (30) is valid for

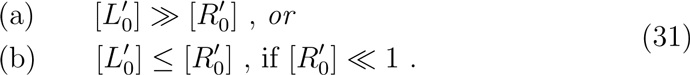

Interestingly, we have a new condition for the use of the PFO approximation in ligand–receptor binding with 1:1 stoichiometry. Equation (31a) is the constraint (12) already in widespread use. However, equation (31b) extends the range of conditions under which PFO dynamics may be applied. The regions of validity of the PFO approximation are illustrated graphically in Fig. 2 by plotting conditions (30) and (31) in the space of initial ligand concentrations, [*L*_0_], and equilibrium constant, *K*_*S*_, normalized by [*R*_0_] Typically, PFO kinetics are assumed valid when the ratio of [*L*_0_]:[*R*_0_] is greater than 10:1 [20, 3, 27] Applying this same “rule of thumb”, we set the left-hand side of (30) equal to 0.1 to separate valid from non-valid regions in Fig. 2. Note that the plane is divided into three regions by lines corresponding to condition (30) and the generally used condition (12). Region B comprises a portion of the space where PFO kinetics were previously assumed to be invalid, but in fact the errors introduced by the approximation in this region are expected to be small, even for initial conditions such that [*L*_0_] ≈ [*R*_0_].

**Figure 2:**
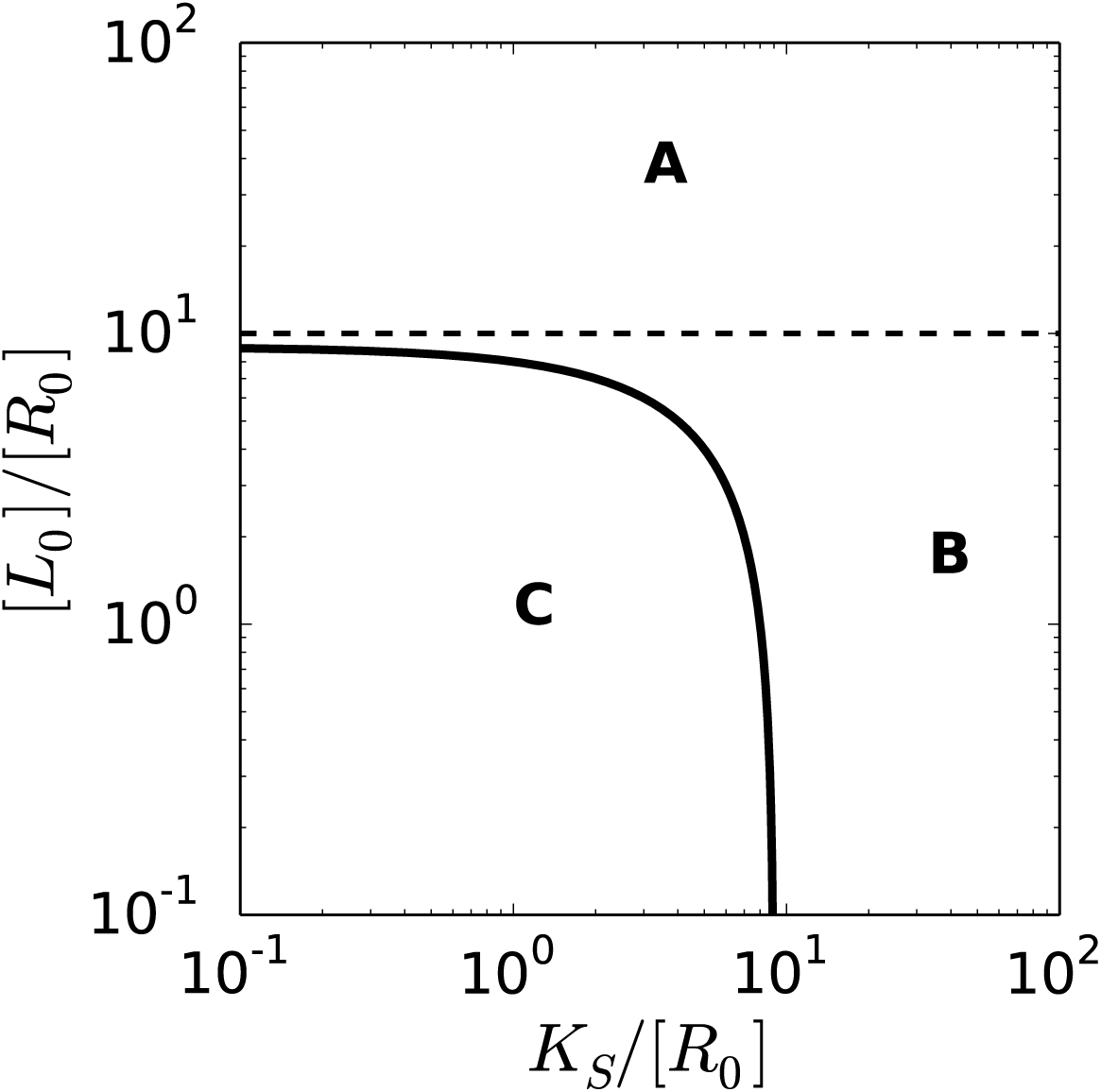
Validity regions in the [*L*_0_]/[*R*_0_]–*K*_S_/[*R*_0_]log-log plane for the use of pseudo-first-order kinetics to model ligand receptor reaction (1). The dashed line indicates 10[*R*_0_] = [*L*_0_], the lower line is 9[*R*_0_] = [*L*_0_] + *K*_*S*_. In region A, [*R*_0_] ≫ [*L*_0_], where PFO kinetics has here been shown to be valid, as previously thought. In region B, [*R*_0_] does not greatly exceed [*L*_0_], but *K*_*S*_ ≫ [*R*_0_]. Here we have shown that PFO kinetics holds, even for [*L*_0_] ≈ [*R*_0_]. In region C, where [*R*_0_] does not greatly exceed [*L*_0_] and *K*_s_/[*R*_0_] is not much greater than 1, PFO kinetics is not valid.

## 5. There are two types of errors observed with the application of approximations in reaction kinetics

In reaction kinetics, there are two type of errors that can be committed when applying an approximation to the governing equations of a complex reaction. Research in mathematical chemistry and biology primarily focuses on estimates of errors in the concentrations of reacting and product species. This error – which we name *concentration error* – is commonly evaluated by calculating the difference between the solution of the approximate equation (16) with that of the exact equation (10) (see, for example, [28]). The concentration error provides a measure of how closely the approximate solution matches the exact solution. However, experimentally, PFO approximations are often used to derive expressions that facilitate estimating kinetic constants using nonlinear regression methods. Therefore, of particular utility are error estimates from fitting kinetic parameters using the mathematical expression derived with the PFO approximation. We called these *estimation errors*.

Naively, it would seem that if the difference between the complex concentration, *C*(*t*), is small between the PFO approximation and exact solution, then the PFO equation should provide accurate estimates of the rate constants when used to model experimental data. This, however, is not necessarily true in general. To better understand why the concentration and estimation errors are not the same, and to show where they diverge, we compare errors based on the numerical difference between the exact and PFO models with those based on estimating rates constants using the PFO model.

### 5.1 Analysis of the concentration error

Theoretically we define a concentration error measure as

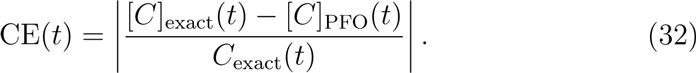

For the bimolecular ligand–receptor binding (1), we can calculate analytically the above expression by replacing [*C*]_exac_(*t*) with (10), and [*C*]_PFO_(*t*) with (16). However, this expression is too cumbersome, and hence we present a numerical analysis of the concentration error. Fig. 3 presents the time course of the exact and approximate complex concentration [Fig. 3(a)–(c)], and a calculation of the percentage concentration error (100 × CE) [Fig. 3(d)–(f)] introduced by the PFO approximation for initial conditions lying in region A, B and C of Fig. 2, respectively. The error remains less than 3% over the course of the reaction for the cases satisfying condition (30), and approaches 30% for the point in region C.

**Figure 3:**
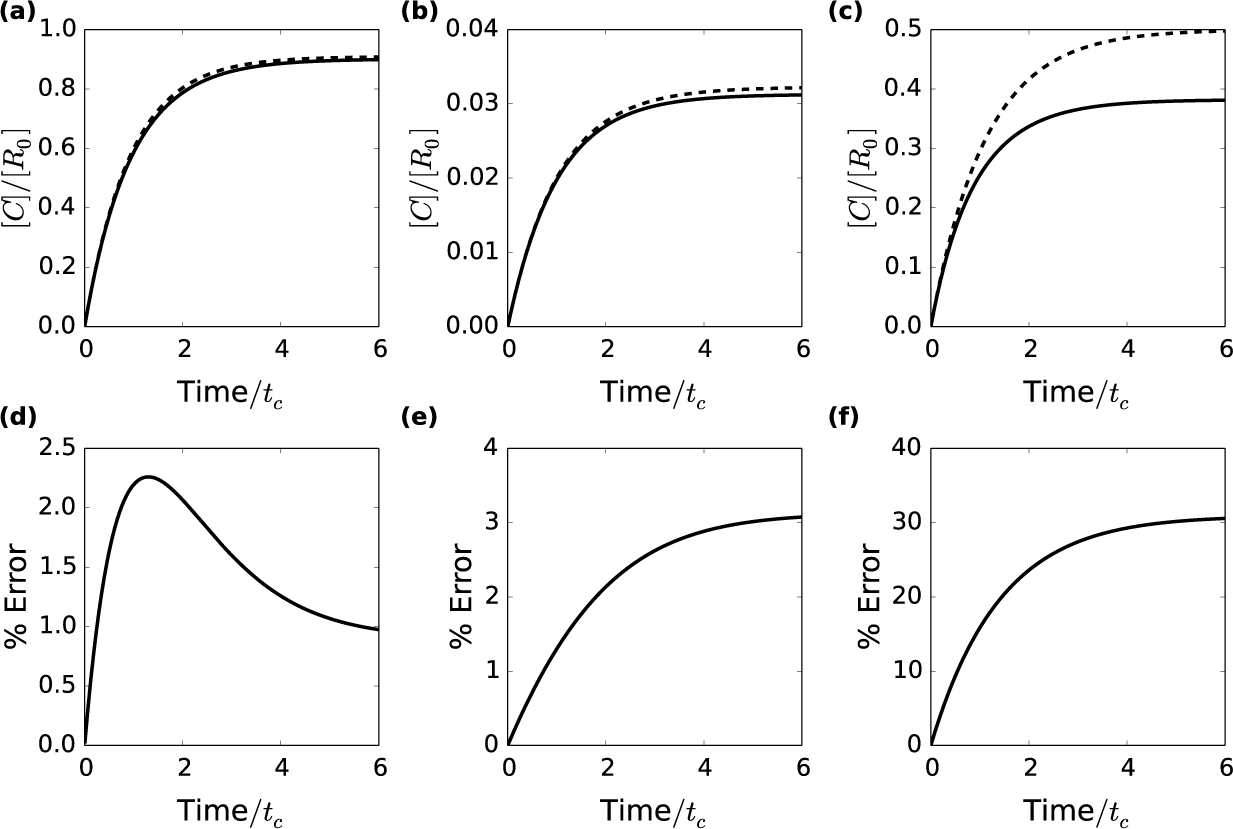
Time course of species concentrations and concentration errors of pseudo-first-order approximation for the ligand receptor reaction (1). Panels (a)-(c) show the concentration of the complex as a function of time for the cases: (a) [*L*_0_] = 10[*R*_0_] and *K*_*S*_ = [*R*_0_], (b) [*L*_0_] = [*R*_0_] and *K*_*S*_ = 30[*R*_0_], (c) [*L*_0_] = [*R*_0_] and *K*_*S*_ = [*R*_0_]. The dashed lines correspond to calculations assuming pseudo-first-order kinetics, while solid lines are exact solutions, (d)-(f) show the errors induced by assuming pseudo-first-order kinetics for case (a)-(c), respectively.

It is useful to define a scalar measure based on (32). For this there are many options, yet in order to remain as conservative as possible, we choose the maximum value of concentration error over the time course of the reaction, which we call the *maximum concentration error*. Additionally, we calculate the *steady-state concentration error*, defined as lim_t→∞_ CE (*t*). Fig. 4 shows the maximum and steady-state concentration error contours for initial conditions in the [*L*_0_]-*K*_*S*_ plane. In general, the maximum concentration error is well-described by the newly-derived condition. The error contours allow for a quantification of what is meant by “much less than”. The commonly used requirement that [*L*_0_] ≥ 10[*R*_0_] produces errors always less than 5%. In fact, when *K*_*S*_ is less than [*R*_0_], a ratio of ligand to receptor of approximately 4:1 is enough to constrain the error to below 5%. Although the condition for the validity is symmetric in [*L*_0_] and [*K*_*S*_], the errors are not. The ratio of *K*_*S*_:[*R*_0_] must be approximately 20:1 for the error to remain below 5%. This is not surprising since the exact solution is not symmetric in [*L*_0_] and *K*_*S*_, so we should expect that the two quantities would have different effects. Nevertheless, the notion that PFO kinetics can rightly be assumed even when the ligand is not present in excess holds true.

**Figure 4:**
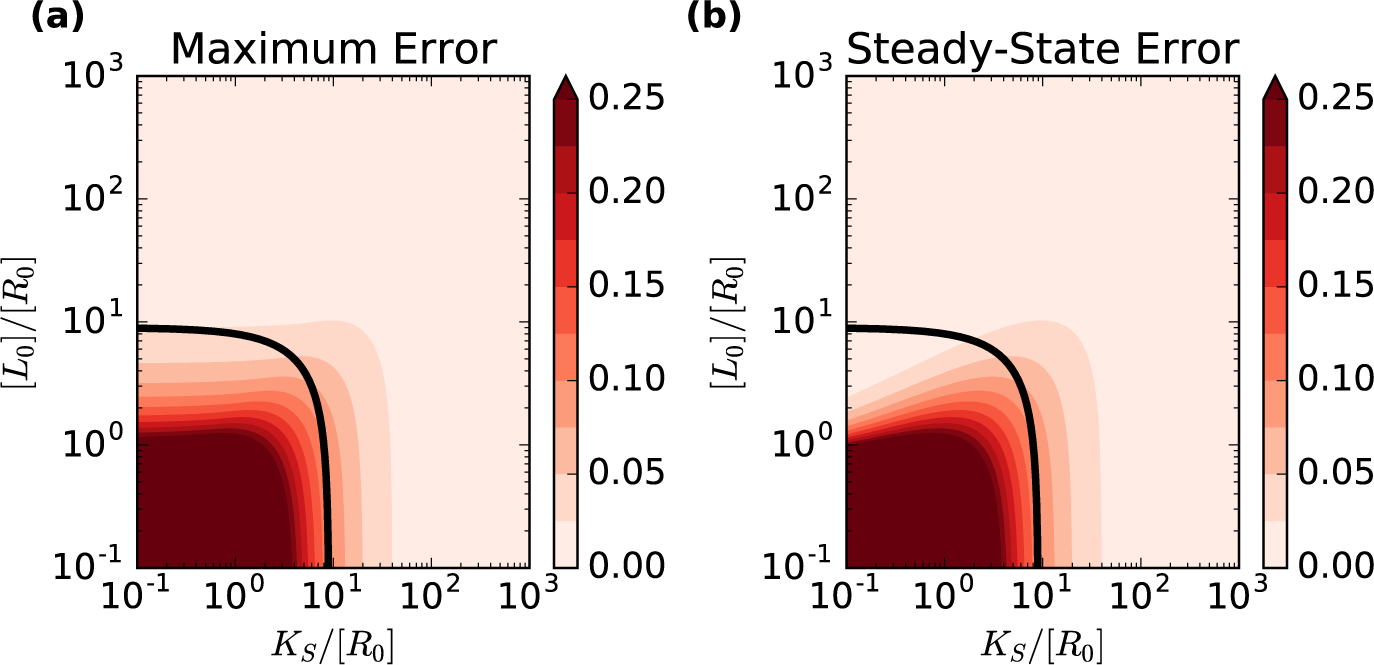
Maximum and steady-state concentration errors for the ligand receptor reaction (1). Panel (a) shows a heat map of the maximum concentration error incurred by assuming pseudo-first-order approximation for different initial conditions. Coloring corresponds to the error as defined in (32). Similarly, Panel (b) shows the steady-state concentration error between the exact and pseudo-first-order solutions. The black lines correspond to condition (30) when the left-hand side is equal to 10.

### 5.2 Analysis of the estimation error

Next, we calculate the error in estimated rate constants by generating sample data using the exact solution. The frequency of the sampling, *ω*_obs_, and the time span of the sampling window, *t*_obs_, are varied. Values of *ω*_obs_ range from 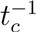 to 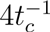. Higher sampling frequencies were also tested, but results are not presented as they did not show appreciable difference from the case of *ω*_*obs*_ = 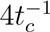. The sampling windows tested begin at *t* = 0 and continue for *t*_*obs*_ = 3*t*_*c*_, 10*t*_*c*_, 100*t*_*c*_. An “experimental protocol” for a numerical experiment then consists of choosing specific *ω*_*obs*_ and *t*_obs_, and using (10) to calculate [*C*] (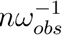) for integers *n* ∈ [0,*t*_obs_*ω*_*obs*_]. For each simulated data set (for which there is no experimental error), the rate constants *k*_*1*_ and *k*_*−1*_ are estimated by fitting the data with equation (16) using the Levenberg–Marquardt algorithm as implemented in SciPy (version 0.17.0, http://www.scipy.org) with initial estimates for *k*_1_ and *k*_−1_ equal to the values used to generate the data. We then define the *estimation error* of a parameter *k*_*i*_ as

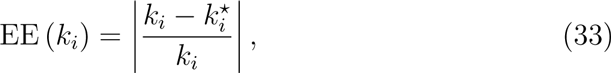

where 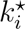 is the estimate of *k*_*i*_ calculated from fitting the PFO solution to the generated data. Additionally, for the ligand–receptor interaction, we calculate the *mean estimation error* as an aggregate measure of the parameter estimation

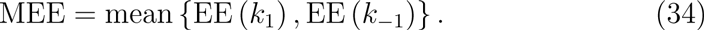

Concentration error measures, such as the maximum or steady-state concentration errors, that compare exact and approximate solutions to the ligand–receptor complex concentration, are fundamentally different from those incurred by using an approximate model to fit experimental data. To illustrate this point, Fig. 5 shows contours for the mean estimation error when different “experimental protocols” are used to generate data. For a given sampling frequency, the mean estimation error contours increasing conform to the contour derived from condition (30) as *t*_*obs*_ increases. However, for initial conditions with [*L*_0_] ≥ 10[*R*_0_] and *K*_*S*_ ≤ [*R*_0_], significant estimation errors can occur if the observation time is not sufficiently long. Even after 10*t*_*c*_ of observation, at which point [*C*](*t*) > 0.999[*C*]*, the mean error in the estimated parameters can exceed 10% when *K*_*S*_ is small. Only after nearly 100*t*_*c*_ the mean estimation error contours closely mimic the theoretical condition for the validity of the PFO kinetics. This highlights the difference between the errors calculated by comparing the exact and approximate solutions of the concentration equations, and those errors due to fitting an approximate model to data. Additionally, it should caution experimentalists from applying PFO approximations whenever one species is in excess. The values of the rate constants must be considered as well.

**Figure 5:**
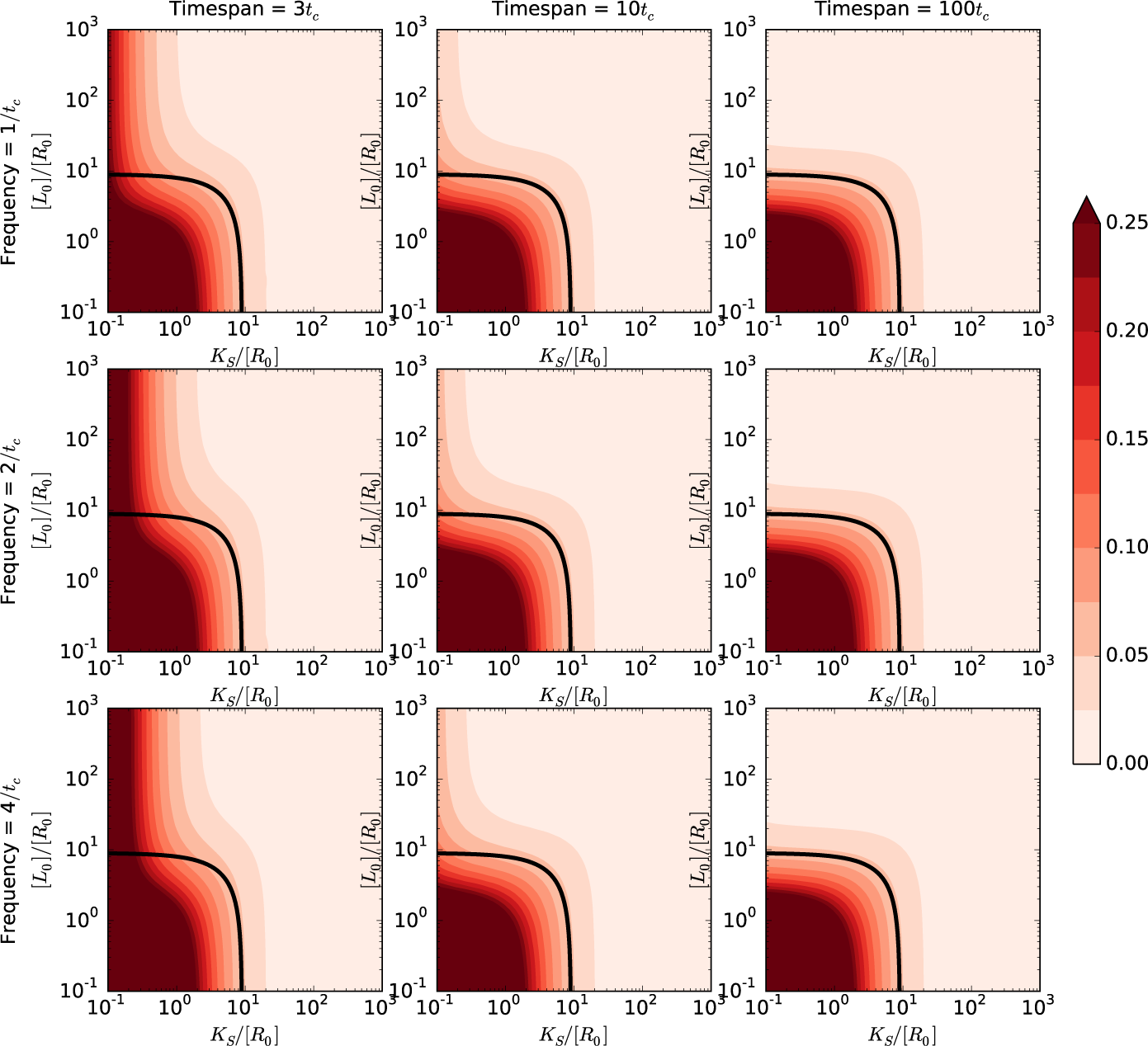
Mean estimation error in the rate constants when applying the pseudo-first-order approximation to the ligand receptor reaction (1). Heat maps of the mean estimation error of the rate constants are shown for different experimental protocols. Columns, from left to right, correspond to increasing length of observation time. Rows, from top to bottom, correspond to increasing frequency of sampling. For short observation windows, large errors occur even when [*L*_0_]≫ [*R*_0_]. Also, counter-intuitively, the error at some initial conditions increases as the sampling frequency increases (e.g. down column 1). The black lines correspond to condition (30) when the left-hand side is equal to 10.

Interestingly, increasing the frequency of sampling does not necessarily improve the estimation of the rate constants. In fact, the first column in Fig. 5 shows, that the mean estimation error actually increases as more sample points are used. This effect saturates quickly as the frequency is increased, but nevertheless, using fewer *exact* data points can lead to improved predictions. One major benefit of numerous data points is that it reduces error due to measurement noise, and this likely will outweigh the gains from using fewer data points when fitting. Yet, in cases where accurate measurements are possible, fitting more data to an approximate model can have deleterious effects on the accuracy of parameter estimates made from such a fit. It may be possible to take advantage of both of these effects by recording data at a high frequency, say using an optical assay [29], then performing a number of fits on subsets of the data sampled at lower frequency, thereby reducing both the experimental noise and the estimation error incurred by fitting to an approximate model.

### 5.3 The error in the estimated parameters for high ligand concentration is due to inaccuracies in k_−1_

Greater understanding of the estimation errors found at high ligand concentrations and low *K*_*S*_ can be gained by comparing the estimation errors of *k*_1_ and *k*_−1_ individually. Fig. 6 shows contours of EE(*k*_1_) [panel (a)] and EE(*k*_−1_) [panel (b)] for numerical data generated with *ω*_*obs*_ = 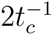 and *t*_*obs*_ = 3*t*_*c*_. From this, it is clear that the inaccuracies lie in predictions of the dissociation rate constant *k*_−1_. The reason *k*_−1_ errors are large for cases in which [*L*_0_] ≫ [*R*_0_] and *K*_*S*_ ≪ [*R*_0_] can be understood by examining the PFO solution (16) in the limit *K*_*S*_/[*L*_0_] → 0. Keeping terms up to linear order in
*K*_*S*_/[*L*_0_], we obtain

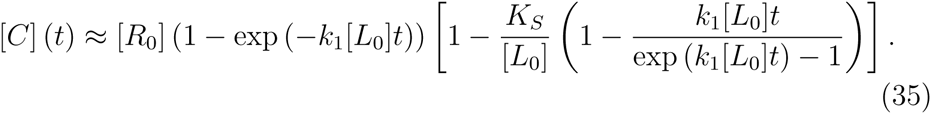

**Figure 6:**
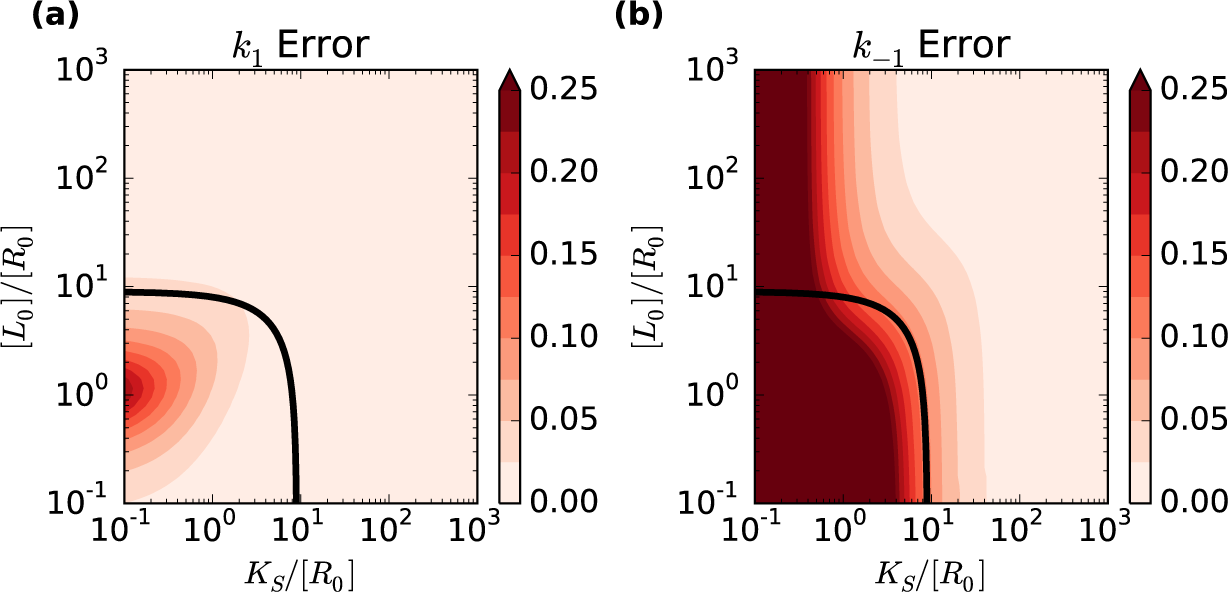
Comparison of estimation errors for *k*_1_ and *k*_−1_. Heat maps of the EE(*k*_1_) and EE(*k*_−1_) are presented in (a) and (b), respectively. The sampling frequency used to generate data was *ω*_*obs*_, = 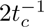 and the observation window was *t*_*obs*_ = 3*t*_*c*_. The black lines correspond to condition (30) when the left-hand side is equal to 10. The error contour for *k*_−1_ estimation shows clear deviations from the analytical conditions, whereas *k*_1_ estimations are accurate where PFO kinetics is shown to be theoretically valid.

The zeroth-order term has no dependence on *k*_−1_. Hence, in this limit there is no unique mapping between the parameters ([*R*_0_], [*L*_0_],*k*_1_, *k*_−1_) and time onto the concentration [*C*], since [*C*] is completely described by the parameters ([*R*_0_], [*L*_0_], *k*_1_) and time. This implies that estimates of *k*_−1_ from progress curve experiments performed under conditions of high ligand concentration and low disassociation constant are unreliable. However, estimates of the association rate constant *k*_1_ from such experiments should be valid. Unfortunately, since *K*_*S*_ is not generally known a priori, a different measurement, such as an equilibrium binding assay, is required to estimate its value. Then, with knowledge of *K*_*S*_, *k*_−1_ can be calculated.

### 5.4 The pseudo-first-order kinetics can lead to significant estimation errors when the conditions of the pseudo-first-order approximation are valid

Taken together, Fig. 4 and Fig. 5 illustrate an important, yet often overlooked, distinction between methods by which to assess the validity of an approximation in chemical kinetics. The first method, popular among theorist, attempts to answer the following question: Given a set of known parameters, how well does the approximate model represent the exact model& This comparison can be made by calculating a maximum or steady-state concentration error, as we do here, or through other measures such as a mean-squared difference over the time course. The second method, which is of greatest importance to experimentalists, answers a subtly different question: Given data, how well do parameters estimated by fitting the data with an approximate model represent the actual parameters& This distinction, between a forward and an inverse problem, is not generally considered when deriving conditions for the validity of reduced kinetic models [30]. To illustrate this, Fig. 7 shows data points generated with exact solution to the governing equations of the ligand–receptor reaction (1), the PFO solution using the same rate constants as those used to generate the exact data points, and the PFO solution using rate constants calculated through nonlinear regression of the simulated data. The PFO solution using the exact rate constants captures the kinetics much more closely than the PFO solution using estimated rate constants, especially as steady-state is approached. Often, it is the former case that is used by theorists to determine valid ranges for an approximation, while the latter case is where the approximation is actually used to interpret data. As we have shown, these two cases are distinct. Hence, when providing ranges for the validity of a simplifying approximations to theory, it is crucial that the application of that theory be kept in mind.

**Figure 7:**
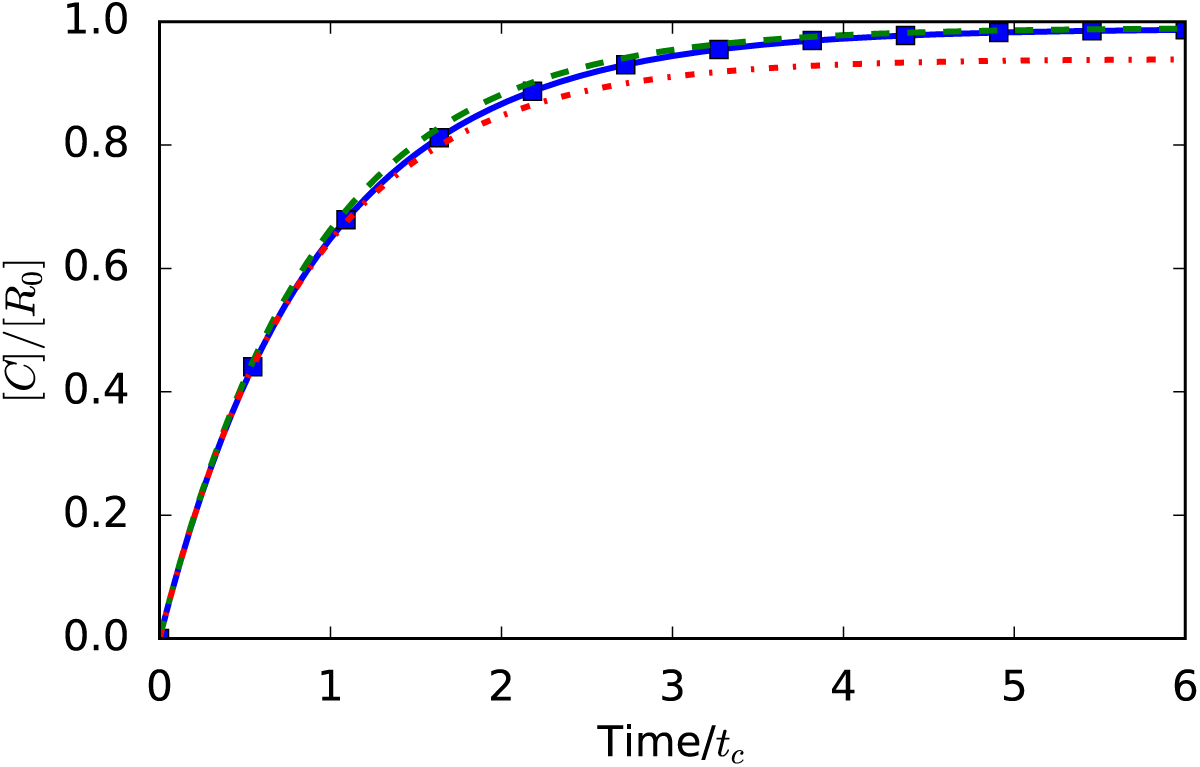
Comparison of approximate model with exact and fitted parameters for the ligand–receptor reaction (1). The blue line represents the exact solution and blue squares are simulated data points. The green dashed line is the pseudo-first-order approximation using the same rate constants used to in the exact solution. The red dot-dashed line is the pseudo-first-order approximation using rate constants found by fitting the pseudo-first-order model to the data generated using the exact solution.

## 6. Discussion

We have investigated the application of the PFO approximation to ligand–receptor binding dynamics. PFO kinetics are used to linearize the solutions to the differential equations that describe the concentration of ligand, receptor and ligand–receptor complex over time, allowing them to be fit by a single exponential [16, 15]. This approximation is known to introduce errors that are acceptably small under certain conditions, which have generally been described by [*L*_0_] ≫ [*R*_0_]. In this paper, we show that this condition is somewhat more stringent than necessary, specifically when [*R*_0_]≪ *K*_*S*_. In fact the condition [*R*_0_] ≪ *K*_*S*_ provides another sufficient condition under which one may safely use the PFO approximation, with little regard to the concentration [*L*_0_].

Although it is possible to derive closed-form solutions that describe the kinetics of simple ligand–receptor binding [see, equation (10)], this equation is cumbersome. The PFO approximation gives a much simpler solution [see, equation (16)], which can be linearized by use of logarithmic plots to facilitate data fitting [17]. With this simpler form a linear fit suffices to determine all of the relevant rate constants leading to a complete description of the reaction kinetics. The new condition developed here extends the validity of this method into new territory, increasing its usefulness. Specifically, in cases where reagents are either expensive or difficult to isolate in significant quantities, the new condition suggests far more economical usage (in some cases, orders of magnitude lower concentration) of reagents is possible.

Additionally, we have shown that there is often an inconsistency between the derivation of conditions for validity of an approximation, and the relevant measure of error for applications of that approximation. Approximations to a theory are generally taken as valid if, using identical input parameters, exact and approximate solutions for species concentration are sufficiently similar. This requires minimizing the concentration error introduced in equation (32). However, approximate models are often used to estimate kinetic parameters through fitting to experimental data. We have demonstrated that estimation errors may be significant, even for conditions in which the approximate and exact models are nearly identical. This effect is particularly apparent when one reactant is in excess, the disassociation constant is small, and the length of the observation is not sufficiently long.

Commonly, experimental protocols for kinetic binding assays call for measurements to be made until concentrations “plateau” [27]. However, the definition of “plateau” is often left to the judgment of the investigator. Our analysis shows that stopping measurement prematurely can lead to significant errors in the rate constants estimated by such experiments. A more rigorous definition of the necessary experimental time to reach plateau should involve the inherent timescale, *t*_*c*_. The error contours in Fig. 5 show that for many initial conditions, specifically for those with large enough *K*_*S*_, measurements over 3*t*_*c*_ are sufficient. It should be possible, however, to test if the experimental conditions are problematic without prior knowledge of *K*_*S*_, so long as the initial concentrations of receptor is known. If, at steady-state, [*C*] is very near [*R*_0_], then the affinity of the ligand for the receptor is high enough (and *K*_*S*_ will be small enough) to make the value of *k*_−1_ from regression analysis unreliable. Since affinities between ligands and receptors are typically quite high, this will often be the case. Hence this suggest that it is necessary to estimate the equilibrium constant using a separate assay. Then, with knowledge of *K*_*S*_, the rate constants can be unambiguously estimated from kinetic data.

Lastly, we emphasize that the difference between the concentration error and estimation error are not specific to the case of ligand–receptor binding. The problem of estimating parameter values for models is well known in the model calibration field [31], and has also received attention from mathematical and systems biologists [30, 32]. The essential questions are whether or not a parameter in a model actually corresponds to the underlying physical property it is meant to represent, and whether the value of the parameter can be uniquely determined from data. Frequently, the value of the parameter that provides the best fit to data differs from the most accurate assessment of the underlying physical property, estimated through some other means. In the case of ligand–receptor binding, the rate constants estimated using PFO kinetics correspond to a best-fit of experimental data to an approximate model. In many cases, these estimates will not coincide with the actual rate constants for the second-order reaction. In fact, this difference is quite general and future studies should investigate how the validity of approximations in, for example, Michaelis–Menten kinetics, or inhibited ligand–receptor binding might change when their ability to accurately predict parameters from data is considered.

## Appendix A. Symmetry of case with no receptor depletion

The second case referred to in the text (Section 2) applies when the concentration of the receptor [*R*_0_] is much greater than that of the ligand [*L*_0_], which implies that

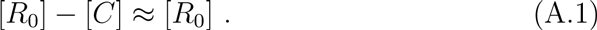

Substituting (36) into equation (7) gives

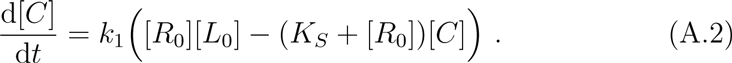

[Compare with (15)]. Solving this equation yields

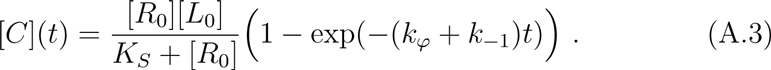

This solution is symmetrical with (16). Condition (36) gives us the following implication parallel with (17)

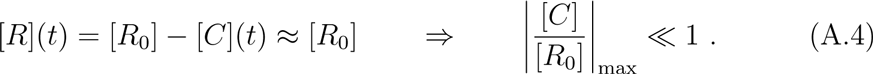

Similar to the case with no ligand depletion, the maximum concentration is equal to λ_. Hence, following the same procedure as in Section 4, a condition, symmetric to (30), for the case with negligible receptor depletion is found to be

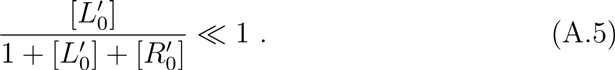

## Appendix B. Validity of approximation (28)

The derivation of the conditions given by (30) requires that equation (28) be satisfied, which we reiterate as

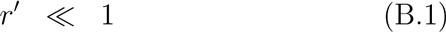

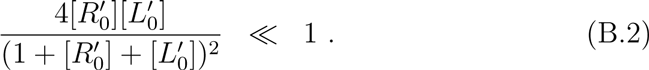

The above inequality can be written as

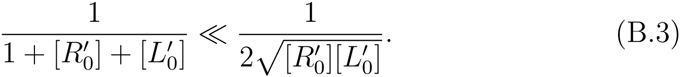

Since the denominators are both positive, we can rearrange this as

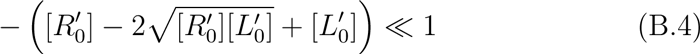

and factoring the left side then gives

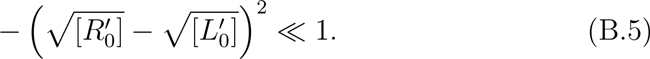

In the above inequalities, the left side is always negative and the right side is clearly positive. Therefore, it is appropriate to assume that 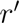 ≪ 1.

## Acknowledgments

This work is supported by the University of Michigan Protein Folding Diseases Initiative.

